# Neural basis of learning to perceive speech through touch using an acoustic-to-vibrotactile speech sensory substitution

**DOI:** 10.1101/2021.10.24.465610

**Authors:** Patrick S. Malone, Silvio P. Eberhardt, Edward T. Auer, Richard Klein, Lynne E. Bernstein, Maximilian Riesenhuber

## Abstract

The goal of sensory substitution is to convey the information transduced by one sensory system through a novel sensory modality. One example is vibrotactile (VT) speech, for which acoustic speech is transformed into vibrotactile patterns. Despite an almost century-long history of studying vibrotactile speech, there has been no study of the neural bases of VT speech learning. We here trained hearing adult participants to recognize VT speech syllables. Using fMRI, we showed that both somatosensory (left post-central gyrus) and auditory (right temporal lobe) regions acquire selectivity for VT speech stimuli following training. The right planum temporale in particular was selective for both VT and auditory speech. EEG source-estimated activity revealed temporal dynamics consistent with direct, low-latency engagement of right temporal lobe following activation of the left post-central gyrus. Our results suggest that VT speech learning achieves integration with the auditory speech system by “piggybacking” onto corresponding auditory speech representations.

**Significance statement:** In sensory substitution, the information conveyed by one sensory system is used to replace the function of another. Blind individuals, for example, can learn to use a visual-to-acoustic sensory substitution device to navigate the world. We tested the hypothesis that sensory substitution can more generally occur in typical individuals, by exploiting the existence of multi-sensory convergence areas in the brain. We trained hearing participants to recognize speech syllables presented as vibrotactile stimulation patterns. Using fMRI and EEG, we show that vibrotactile speech learning integrates with the auditory speech system by “piggybacking” onto corresponding auditory speech representations. These results suggest that the human brain can bootstrap new senses by leveraging one sensory system to process information normally processed by another.

## Introduction

A fundamental goal of sensory systems in the brain (e.g., for vision, hearing or touch) is to process physical signals from the environment. Sensory information gathered by peripheral receptors is transduced to the central nervous system in order to extract meaning from stimuli in the environment and guide behavior. Each sensory system specializes in the processing of certain types of physical signals. With the invention of sensory substitution devices (SSDs, (Bach-y-Rita and Kercel, 2003; Bach-y-Rita et al., 1969; Meijer, 1992; Rauschecker, 1995) however, an idea emerged that the information transduced by one sensory system could be used to replace or compensate for the function of another. SSDs use a combination of software and hardware to transform physical signals ordinarily transduced by one sensory system into signals for another system, often in order to compensate for a specific impaired sensory system. Blind individuals, for example, can be trained to use a visual-to-acoustic SSD such as the vOICe (Meijer, 1992) or a visual-to-tactile SSD such as the BrainPort (Danilov and Tyler, 2005; Ptito et al., 2005) to navigate obstacles in the environment. An impressive example of sensory substitution is known as the *Tadoma* method (Alcorn, 1945). Used by extensively trained deaf-blind individuals, the *Tadoma* method involves placing a “listener’s” hand on a talker’s face: The perceiver’s thumb is placed on the speaker’s lips, the middle three fingers on their cheeks and jawline, and the little finger on their larynx. Through proprioceptive and vibrotactile cues resulting from speech production, the perceiver can learn to understand continuous speech (Alcorn, 1945; Norton et al., 1977; Reed et al., 1985). The *Tadoma* method served as an existence proof that individuals could learn to understand speech through their sense of touch with sufficient training. This demonstration converged with the almost century-long effort (Gault, 1924; 1926a; 1926b; 1927) to develop vibrotactile (VT) speech aids, i.e., SSDs that transform acoustic speech into patterns of vibratory stimuli that are presented to the skin. Interest in these devices peaked around the advent of the cochlear implant (Zeng, 2004) in the 1980s, when the long-term consequences of implantation were unknown, particularly for children, and VT devices were competitive with some cochlear implant outcomes. Decades of research with VT speech aids has demonstrated that both hearing and hearing-impaired individuals can be trained to successfully use these devices (Auer et al., 1998; Bernstein et al., 1991; Brooks and Frost, 1983; Eberhardt et al., 1990; Osberger et al., 1993; Rothenberg and Molitor, 1979; Weisenberger and Percy, 1995; Weisenberger et al., 1991).

What is the neural basis of VT speech learning and perception? One possible mechanism is the cross-modal recruitment of auditory cortical regions for the processing of vibrotactile speech stimuli, thereby allowing vibrotactile speech stimuli to “piggyback” onto and leverage existing auditory speech processing circuits. Interestingly, a large behavioral and neuroimaging literature has demonstrated multisensory and cross-modal interactions between these two systems. At the behavioral level, for instance, tactile stimuli presented simultaneously with auditory tones improve sound detection and perceived loudness (Gillmeister and Eimer, 2007; Schürmann et al., 2004), and inaudible air-puffs presented to the cheek can alter the speech sound heard by a perceiver (Gick and Derrick, 2009). At the neural level, experiments in humans, macaques, and cats have identified several auditory-somatosensory convergence zones (with convergence defined as the confluence of input from different sensory modalities in the same brain area) that may mediate these behavioral effects including in the brainstem (Aitkin et al., 1981; Kanold and Young, 2001; Populin and Yin, 2002), and in the auditory cortex including the superior temporal gyrus (STG) (Auer et al., 2007; Foxe et al., 2002; Schroeder and Foxe, 2002; Schürmann et al., 2006), the posterior superior temporal sulcus (pSTS) (Beauchamp et al., 2008), and the planum temporale (PT) (Auer et al., 2007).

Convergence between the auditory and somatosensory systems appears to be influenced by experience. For example, there is increased activation in the auditory cortex of deaf individuals in response to vibrotactile stimuli derived from speech, and it was hypothesized that the cross-modal recruitment of auditory cortex was due to life-long somatosensory stimulation due to high-power hearing aid use that produces mechanical vibrations in the ear canal (Auer et al., 2007; Bernstein et al., 1998). Cross-modal plasticity, or the recruitment of brain regions typically specialized for the processing of one sensory modality by another (Allman et al., 2009; Bavelier and Neville, 2002; Levänen et al., 1998; Meredith and Lomber, 2011; Rauschecker and Harris, 1983; Rauschecker and Korte, 1993; Rauschecker and Kniepert, 1994; Rauschecker et al., 1992), is a well-known effect of sensory loss. Whether cross-modal recruitment occurs in healthy adults with intact sensory systems is controversial (Amedi et al., 2007; Kim and Zatorre, 2011; Kupers et al., 2006; Ptito et al., 2005; Saito et al., 2006; Siuda-Krzywicka et al., 2016). Some studies have provided evidence that training with SSDs (to substitute a “novel” for a “familiar” sense, e.g., novel tactile input for familiar auditory input) leads to the cross-modal engagement of brain areas specialized for the familiar sense to process input from the novel sense (Amedi et al., 2007). It is unclear, however, whether the cross-modal engagement of these areas proceeds in a bottom-up or top-down manner. In other words, does cross-modal processing reflect the direct, low-latency engagement of sensory hierarchies in the canonical sense for the processing of input from the novel sense (i.e., the aforementioned “piggybacking” of the novel sense onto circuits specialized for the processing of the canonical sense), or the top-down activation of these brain areas after the stimulus has already been processed, akin to the effects of mental imagery of speech (Tian et al., 2016)?

In the present study, we conducted fMRI and EEG experiments to determine if and how SSDs functionally integrate into existing native sensory processing hierarchies. Specifically, we investigated whether experience with a vibrotactile speech aid results in the cross-modal recruitment (i.e., engagement) of the auditory system. We trained normal-hearing individuals to identify nonsense vowel-consonant-vowel (e.g., /aba/) disyllables that were presented on an MRI-compatible custom-built 14-channel VT display. Representational similarity analysis (RSA) (Kriegeskorte and Kievit, 2013) of fMRI data acquired before and after training was used to characterize the neural representation of VT speech and the effects of training. In a separate auditory scan, the representation of acoustic speech was localized to determine if any brain regions were selective for speech transduced through different sensory systems. RSA and source-estimation of EEG data were performed to determine the temporal dynamics of VT speech processing.

Participants were trained in a paradigm designed to discriminate between phonological features of VT speech syllables. The information in consonants and vowels is often expressed in terms of phonological categories or dimensions that are aligned with speech production (Zsiga, 2012). Here, we focused on consonant phonological features, which were voicing (the temporal proximity of vocal fold vibration with supralaryngeal closure, e.g., “b” vs. “p”), manner of articulation (the type and degree of constriction during articulation, e.g., “s” vs. “t”), and place of articulation (specifying the point of constriction in the vocal tract during articulation, e.g., “m” vs. “n”). The training paradigm focused on the discrimination of phonological features, because phonological features capture abstract speech categories that generalize over low-level physical stimulus features. Given the massive non-invariance in speech signals (e.g., due to different vocal tracts, different phonemic contexts, etc.), the ability to extract an invariant representation of speech is critical to speech perception. An additional advantage of training phonological features of VT speech stimuli is that the neural representation of auditory speech has been posited to be organized according to phonological feature dimensions (Mesgarani et al., 2014). Therefore, training phonological feature contrasts of VT speech may encourage the deployment of brain regions involved in the representation of auditory speech.

## Results

### Participants successfully learn to discriminate phonological features of vibrotactile speech

Participants completed an average of 9.4 +/- 0.58 (mean +/- SEM) training sessions (range: 6-15 sessions). Generalized linear mixed-modelling of response accuracy revealed a significant fixed-effect of training session (*p*<0.001), indicating accuracy increased with training. In the model, session and type of phonological feature were modelled as fixed effects and subject and stimulus were modeled as random effects. There was no significant effect of phonological feature contrast on accuracy in the generalized linear-mixed model. The mean accuracy for oddball detection in the pre-training VT scan was 70.8 ± 7.7% (mean +/- SEM) and was 83.5 ± 6.3% in the post-training VT scan. Generalized linear mixed-modelling of time-point (pre-training vs. post-training MRI scan) revealed that accuracy was significantly higher post-training (*p*=0.013). The mean accuracy for oddball detection in the auditory scan was 92.7 ± 2.7%.

### fMRI representational similarity analysis and task-evoked functional connectivity reveal the representation of vibrotactile speech and its integration with the existing auditory speech system

Representational similarity analysis of fMRI data was performed to localize the neural representation for VT and auditory speech. In the VT scan acquired before VT speech training, significant correlations with the phonological feature RDM were localized to a left-lateralized network of regions including the supramarginal gyrus, posterior superior temporal gyrus and sulcus, inferior parietal lobule, and middle temporal gyrus (Fig. 2B). After training, selectivity for VT phonological features was found in the left ventral post-central gyrus, right superior temporal gyrus, right planum temporale, and right inferior frontal gyrus (Fig. 2C; Table 2). In order to determine which regions showed a significant increase in selectivity for VT speech with training, a paired *t-*test between the pre- and post-training RSA maps was performed. There was significantly greater selectivity for VT speech in the left post-central gyrus and left posterior middle temporal gyrus after training (Fig. 2D). All analyses were thresholded at a voxel-wise *p*<0.001 and cluster-level *p*<0.05 (FWE-corrected).

**Figure 1.**
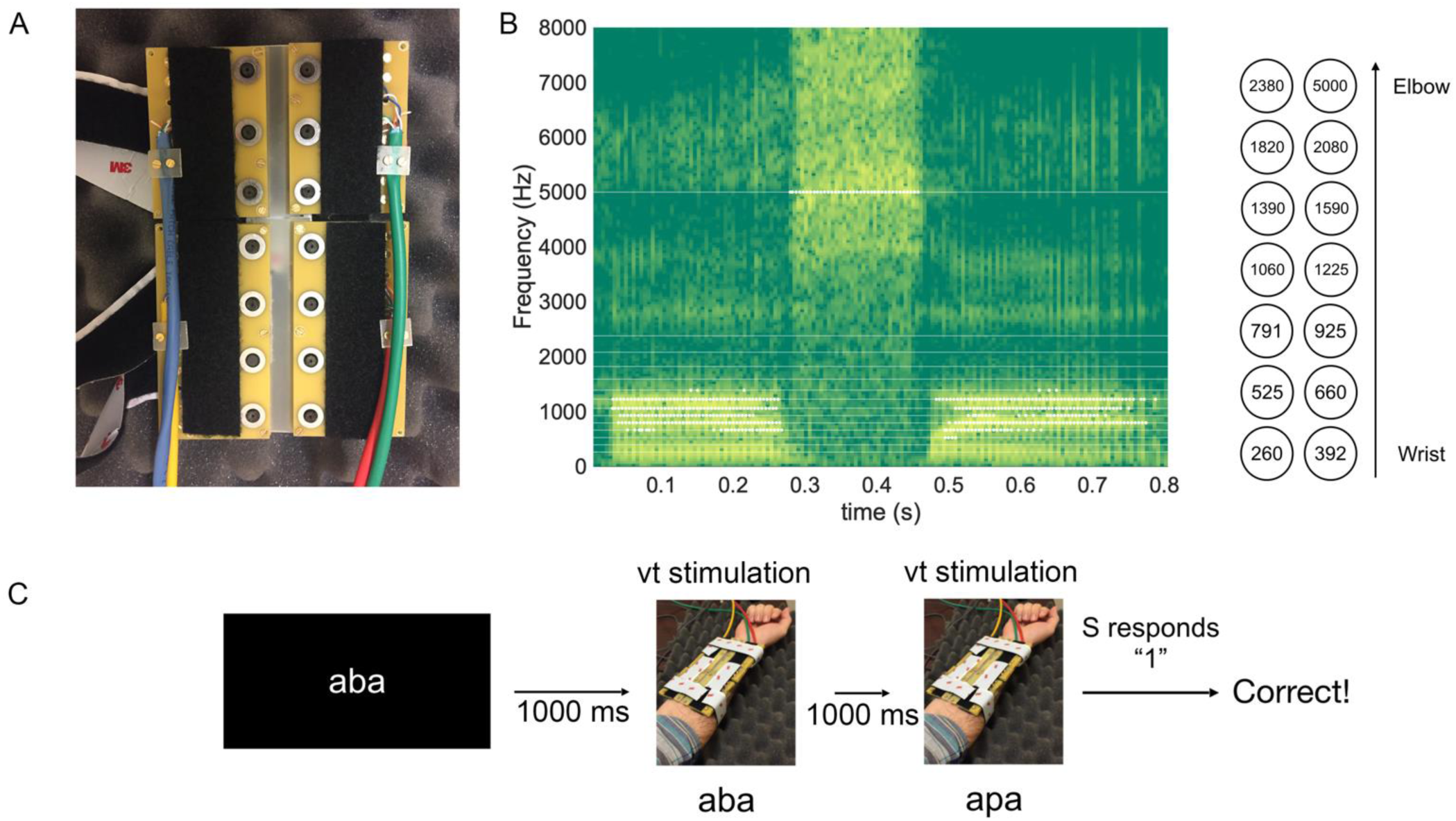
Vibrotactile speech device, stimuli, and training paradigm. **A.** 14- channel vibrotactile speech device. The stimulator contactor side is shown. **B.** Acoustic spectrogram for the disyllable /asa/, overlaid with the corresponding vibrotactile speech stimulus. The horizontal white lines on the spectrogram and the circles to the right of the spectrogram indicate the center frequencies for the vocoder band-pass filters, as well as its high-pass filter at 5000 Hz. The white circles on the spectrogram indicate vibrotactile pulses on the stimulator channels. **C.** Example training trial that trains distinctions in voicing. Participants performed a two-alternative forced-choice paradigm for which the two disyllables differed according to a single phonological feature dimension.

**Figure 2.**
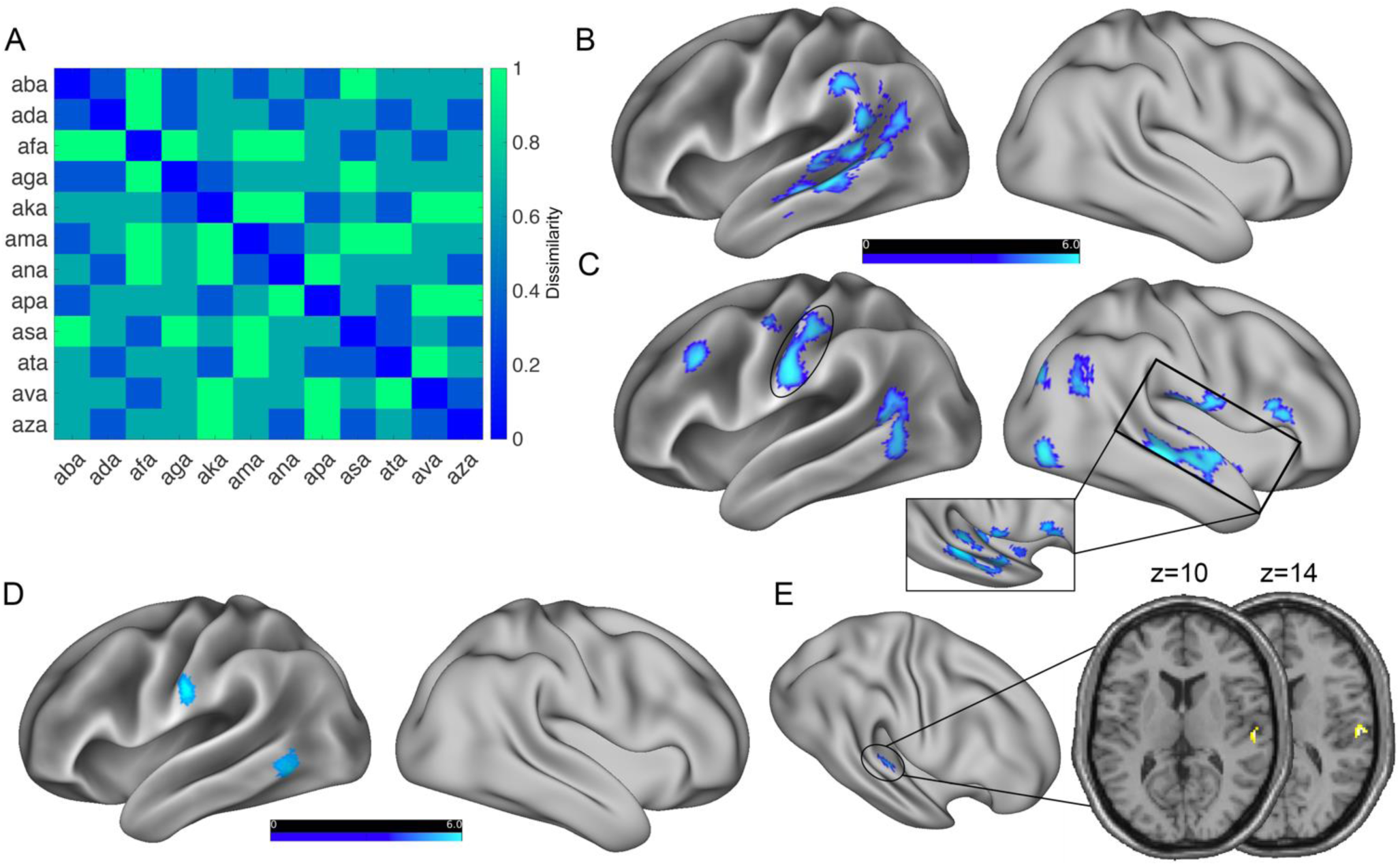
Representational similarity analysis reveals the neural representation of vibrotactile and auditory speech. **A.** Phonological feature representational dissimilarity matrix. **B.** Selectivity for phonological features of vibrotactile speech pre-training. **C.** Selectivity for phonological features of vibrotactile speech post-training. The ellipse indicates the left post-central ROI used as a seed in the functional connectivity analysis in Fig. 3. **D.** Increased selectivity for VT speech with training (paired *t*-test post>pre-training). **E.** Selectivity for phonological features of acoustic speech. Statistical maps in A-D are thresholded at a voxel-wise *p*<0.001 and cluster-level *p*<0.05 (FWE-corrected). Maps in E are thresholded at a voxel-wise *p*<0.001 and cluster-level *p*<0.05 small-volume corrected with the VT speech selective mask in C. Color-bars indicate *t*-statistic.

**Table 1.**
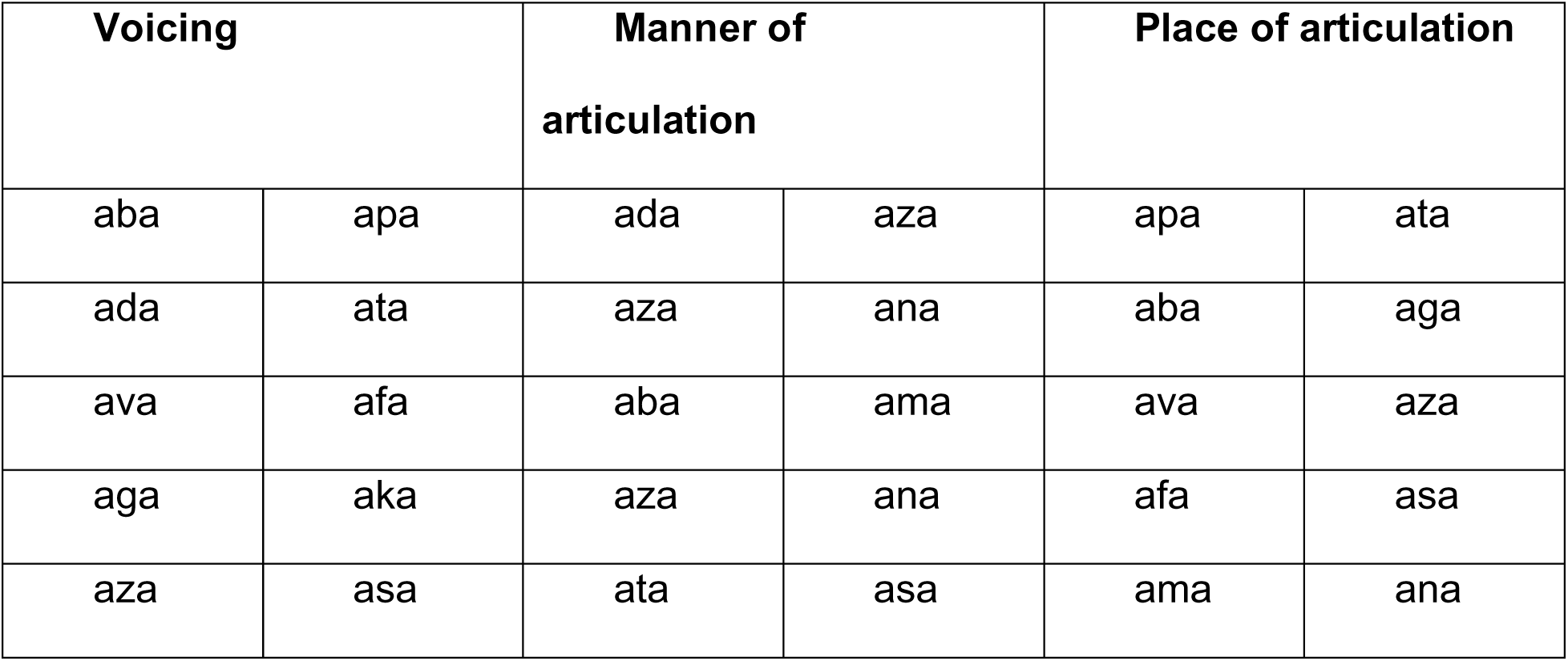
Stimulus pairs used in the vibrotactile speech training task. Each pair of stimuli differed according to one phonological feature (voicing, manner, or place of articulation).

| Voicing |  | Manner of articulation |  | Place of articulation |  |
| --- | --- | --- | --- | --- | --- |
| aba | apa | ada | aza | apa | ata |
| ada | ata | aza | ana | aba | aga |
| ava | afa | aba | ama | ava | aza |
| aga | aka | aza | ana | afa | asa |
| aza | asa | ata | asa | ama | ana |

**Table 2.**
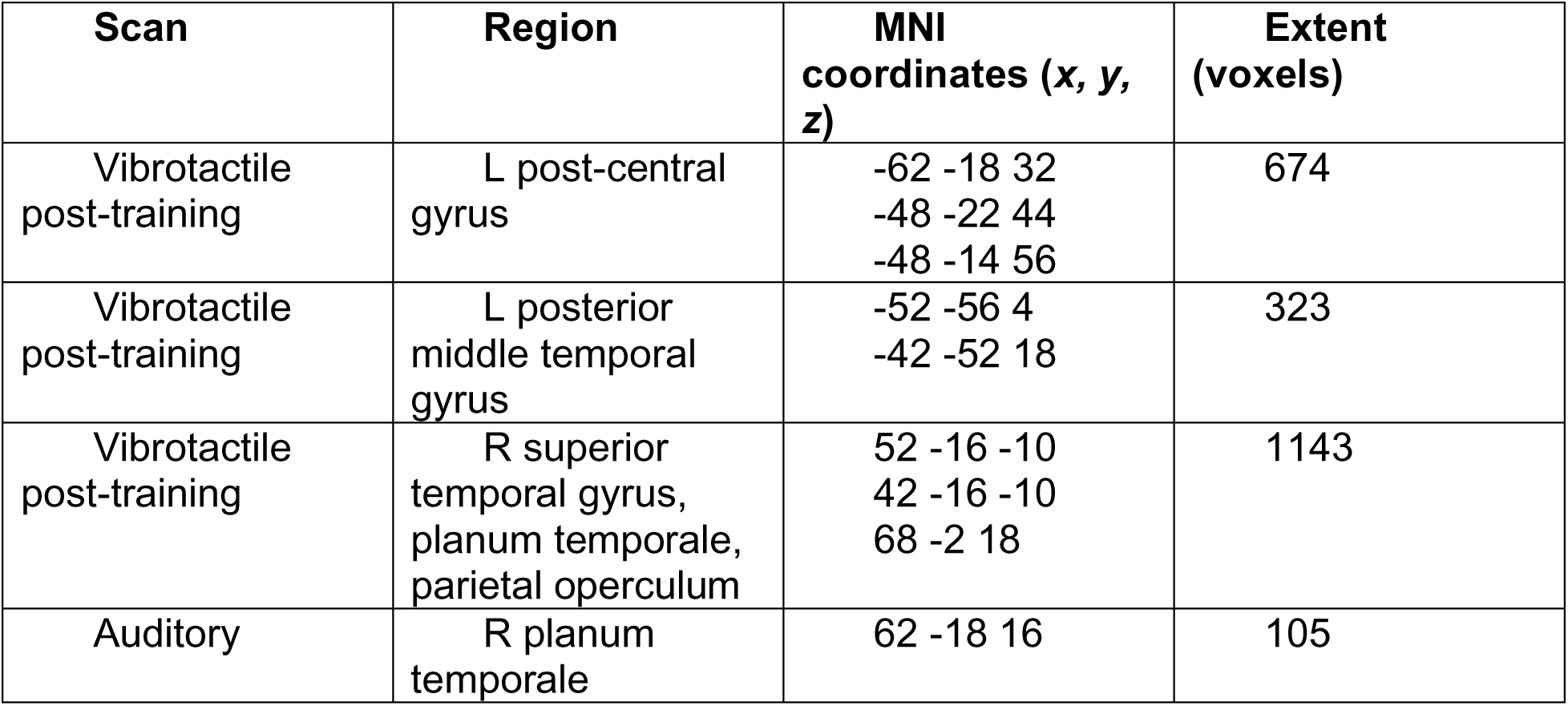
Location and cluster extent for regions-of-interest identified in the fMRI representational similarity analysis.

| Scan | Region | MNI coordinates (x, y, z) | Extent (voxels) |
| --- | --- | --- | --- |
| Vibrotactile post-training | L post-central gyrus | -62 -18 32<br>-48 -22 44<br>-48 -14 56 | 674 |
| Vibrotactile post-training | L posterior middle temporal gyrus | -52 -56 4<br>-42 -52 18 | 323 |
| Vibrotactile post-training | R superior temporal gyrus, planum temporale, parietal operculum | 52 -16 -10<br>42 -16 -10<br>68 -2 18 | 1143 |
| Auditory | R planum temporale | 62 -18 16 | 105 |

In order to test the hypothesis that VT speech learning results in the engagement of brain areas that encode auditory speech, participants underwent an additional post-training fMRI scan in which they listened to the natural (non-vocoded) acoustic versions of the same syllables used in VT speech training. The same RSA as above was performed to localize the representation of auditory speech stimuli. The RSA for the auditory scan was masked by the post-training VT speech selective map (Fig. 2C) in order to isolate regions that were selective for VT speech. Selectivity for phonological features of the acoustic stimuli was identified in the right planum temporale (Fig. 2E; voxel-wise *p*<0.001 and cluster-level *p*<0.05 small-volume corrected with the VT speech-selective mask in Fig. 2C; Table 2).

In order to test the hypothesis that the processing of VT speech involves communication between the somatosensory and auditory systems, seed-to-voxel task-based functional connectivity was performed to characterize the functional connectivity of the left post-central gyrus (seed defined as the left post-central gyrus ROI in Fig. 2C). Pre-training, the left post-central gyrus exhibited functional connectivity with bilateral pre- and post-central gyri, bilateral superior parietal lobule, left planum temporale, bilateral anterior supramarginal gyri, and other regions (Fig. 3A). Post-training, the left post-central gyrus was functionally connected with the aforementioned regions, and new connectivity emerged with bilateral superior temporal gyri, the right PT, and the right middle temporal gyrus (Fig. 3B). Direct comparison of the pre- and post-training functional connectivity maps did not reveal any significant differences in connectivity.

**Figure 3.**
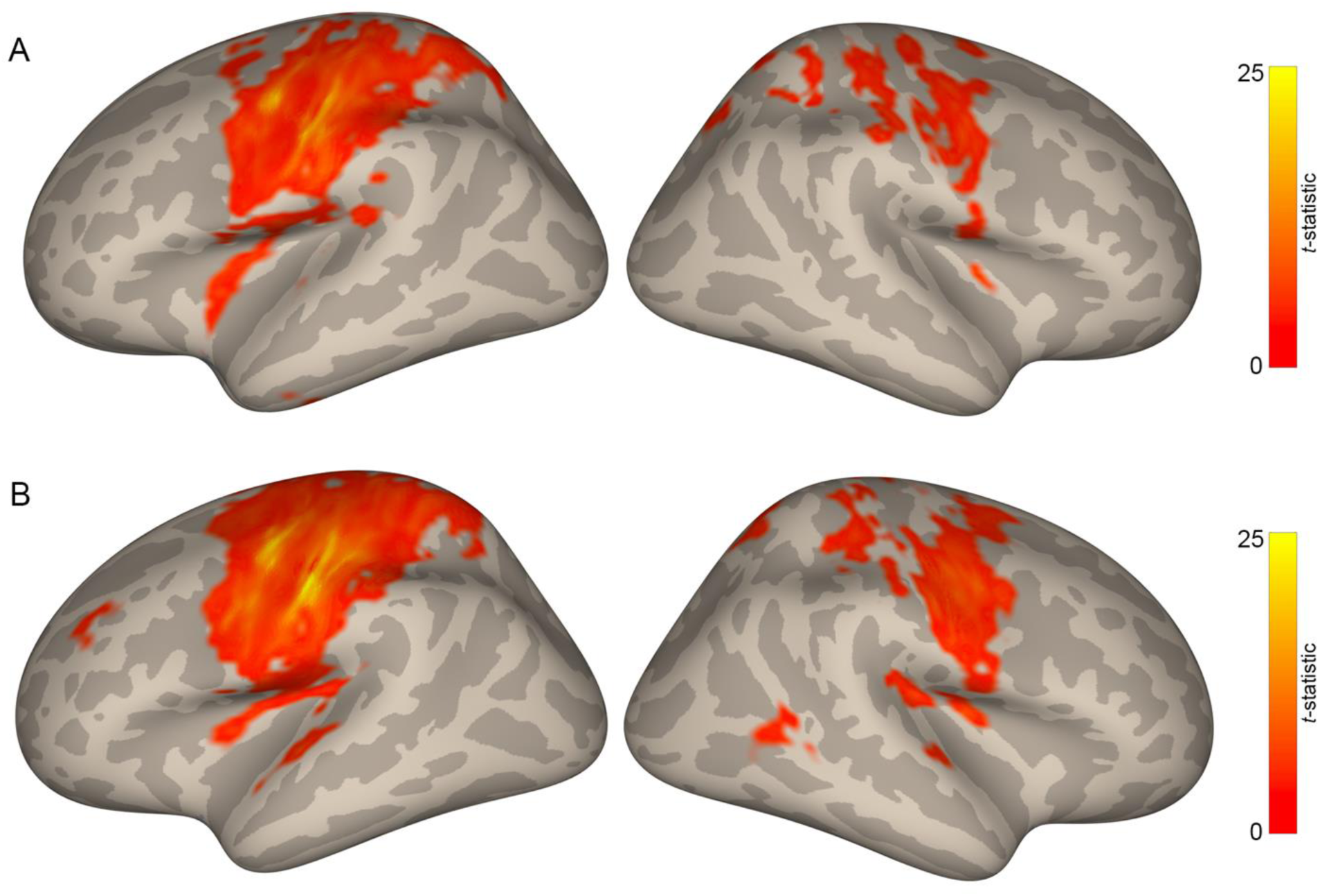
The left post-central gyrus exhibits task-evoked functional connectivity with auditory regions after vibrotactile speech training. **A.** Functional connectivity of left post-central gyrus for the pre-training vibrotactile scan. **B.** Functional connectivity of left post-central gyrus for the post-training vibrotactile scan. The left post-central ROI used as a seed for the connectivity analyses is highlighted with an ellipse in Fig. 2C. All maps are thresholded at a voxel-wise *p*<0.001 and cluster-level *p*<0.05 (FWE-corrected).

### EEG representational similarity analysis and source estimation reveal temporal dynamics of vibrotactile and auditory speech processing

We performed searchlight EEG RSA in order to determine the temporal dynamics of neuronal selectivity for phonological features during the processing of VT and auditory stimuli. Neural RDMs were constructed for each time window and searchlight cluster using the ERPs for auditory and VT stimuli, respectively. For auditory stimuli, neuronal phonological feature selectivity emerged beginning around 330 ms after stimulus onset and lasted until 480 ms after stimulus onset (Fig. 4). For VT stimuli, neuronal phonological feature selectivity began at 480 ms after stimulus onset and lasted until 740 ms after stimulus onset (Fig. 5). The average consonant onset across the 12 VCV syllables was 308 +/- 12.2 ms (mean +/- SEM) after stimulus onset (determined through manual inspection of the acoustic waveforms and spectrograms). Therefore, selectivity for auditory syllables emerged from 28-172 ms after consonant onset, while selectivity for VT syllables emerged 172-432 ms after consonant onset. Interestingly, we observed phonological feature selectivity in right-lateralized channels for both auditory and VT syllables, consistent with our finding of selectivity in the right temporal cortex in the fMRI. The longer latency for VT syllable selectivity could therefore be due to additional signaling latencies from left somatosensory (see Fig. 2D) to right auditory cortex.

**Figure 4.**
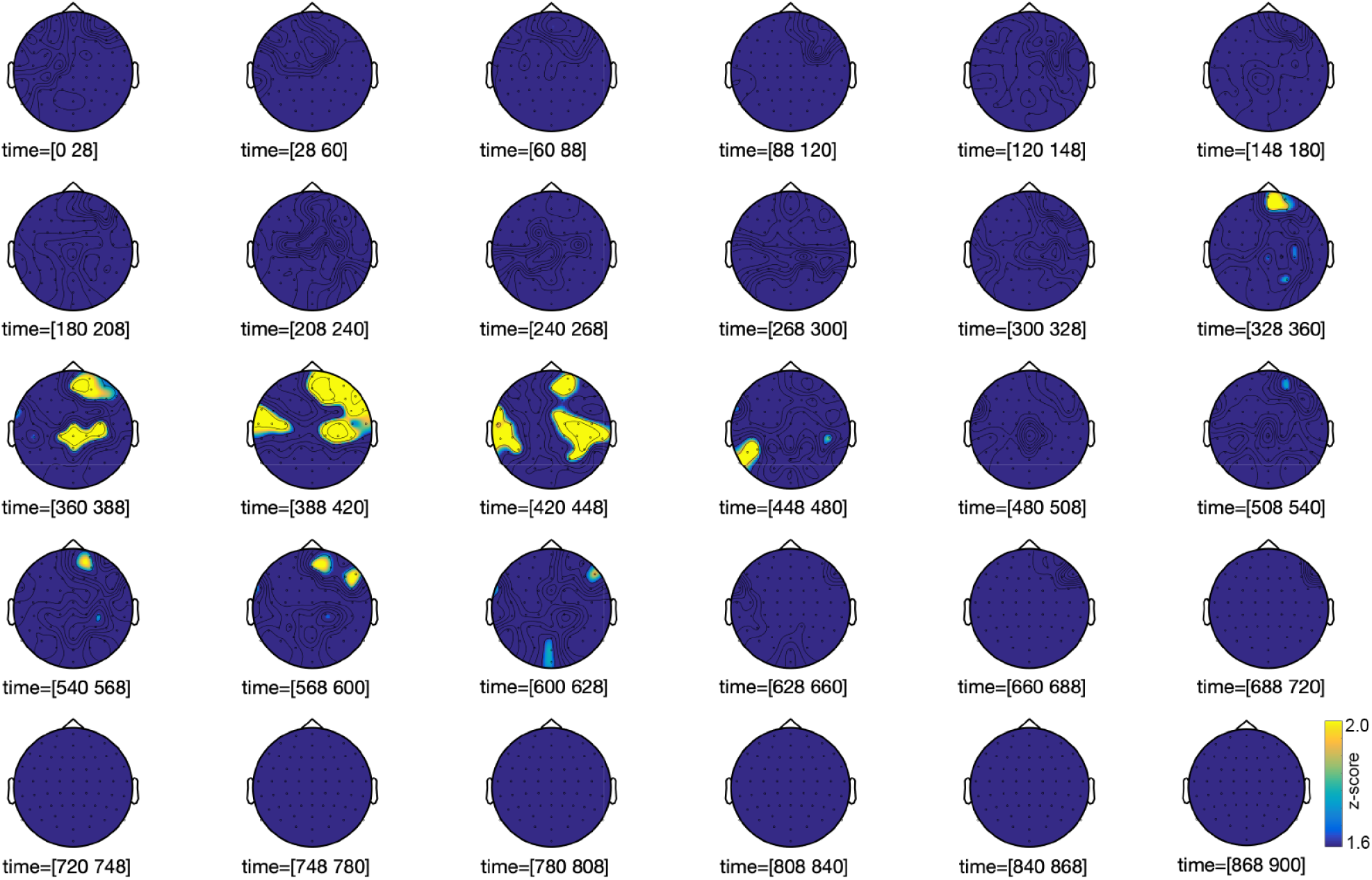
EEG searchlight representational similarity analysis of event-related potentials for auditory stimuli using the phonological feature representational dissimilarity matrix. Topographic plots show z-scores thresholded at *p*<0.05 cluster-level corrected using threshold-free cluster enhancement (see Methods).

**Figure 5.**
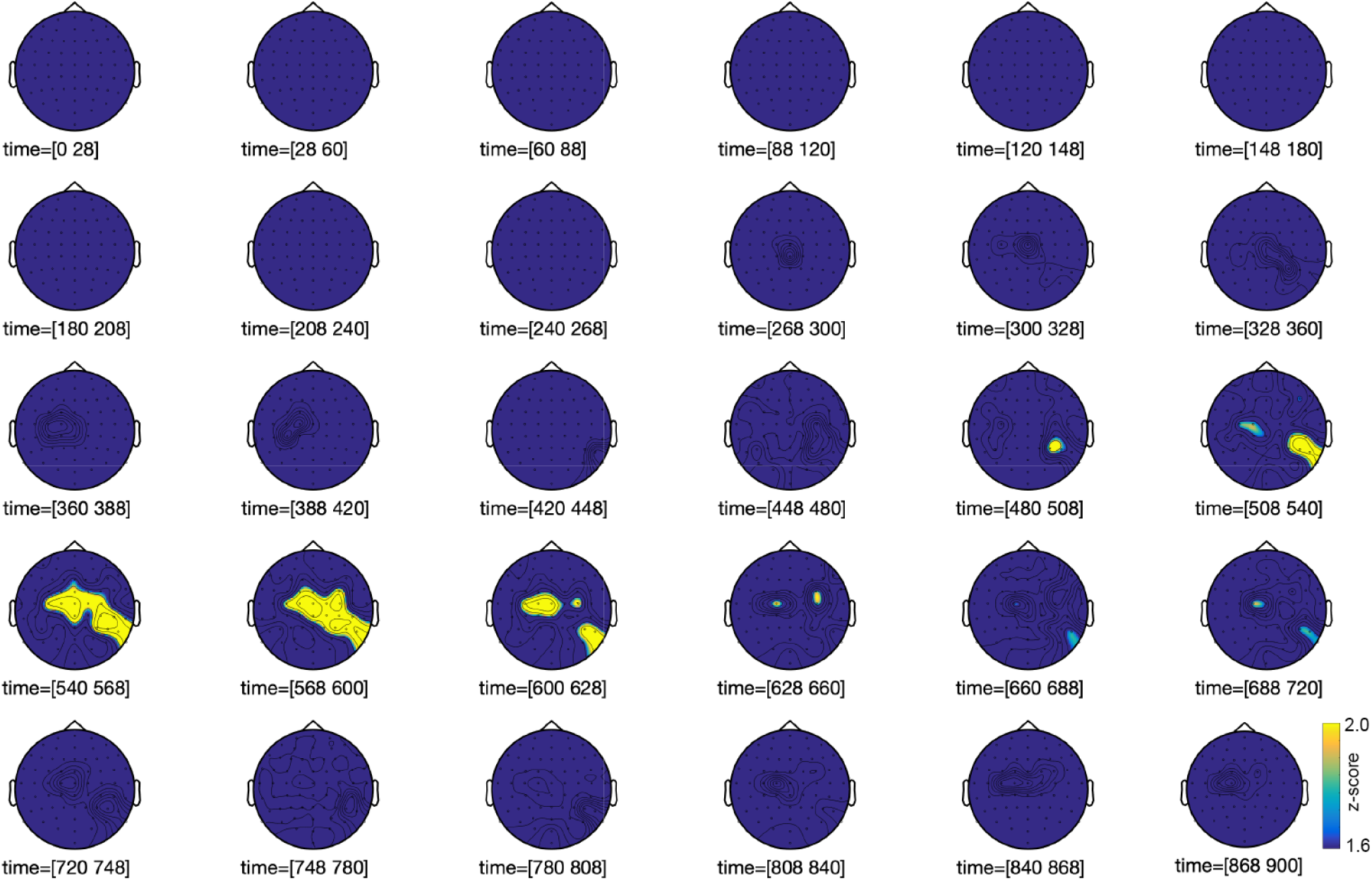
EEG searchlight representational similarity analysis of event-related potentials for vibrotactile stimuli using the phonological feature representational dissimilarity matrix. Topographic plots show z-scores thresholded at *p*<0.05 cluster-level corrected using threshold-free cluster enhancement (see Methods).

In order to determine the temporal dynamics of activation of the two key ROI identified in the fMRI-RSA (the left post-central gyrus and right temporal regions), source estimation of the EEG data was performed. In particular, we were interested in the temporal dynamics of engagement of the right temporal cortex during the processing of VT speech, and whether the latencies were more consistent with bottom-up or top-down processing. During the processing of auditory stimuli, there were early peaks at 48, 100, and 168 ms after stimulus onset in the left post-central gyrus, while activity in right PT/STG peaked at 48 and 128 ms after stimulus onset (Fig. 6B). During the processing of VT stimuli, the left post-central gyrus exhibited two initial peaks in activity at 92 and 144 ms, while the response in right auditory areas peaked at 160 ms (Fig. 6A). Here again, the latency of engagement of the left post-central gyrus earlier than right auditory areas suggests that the VT stimuli were processed first in left somatosensory regions before being relayed to right auditory areas.

**Figure 6.**
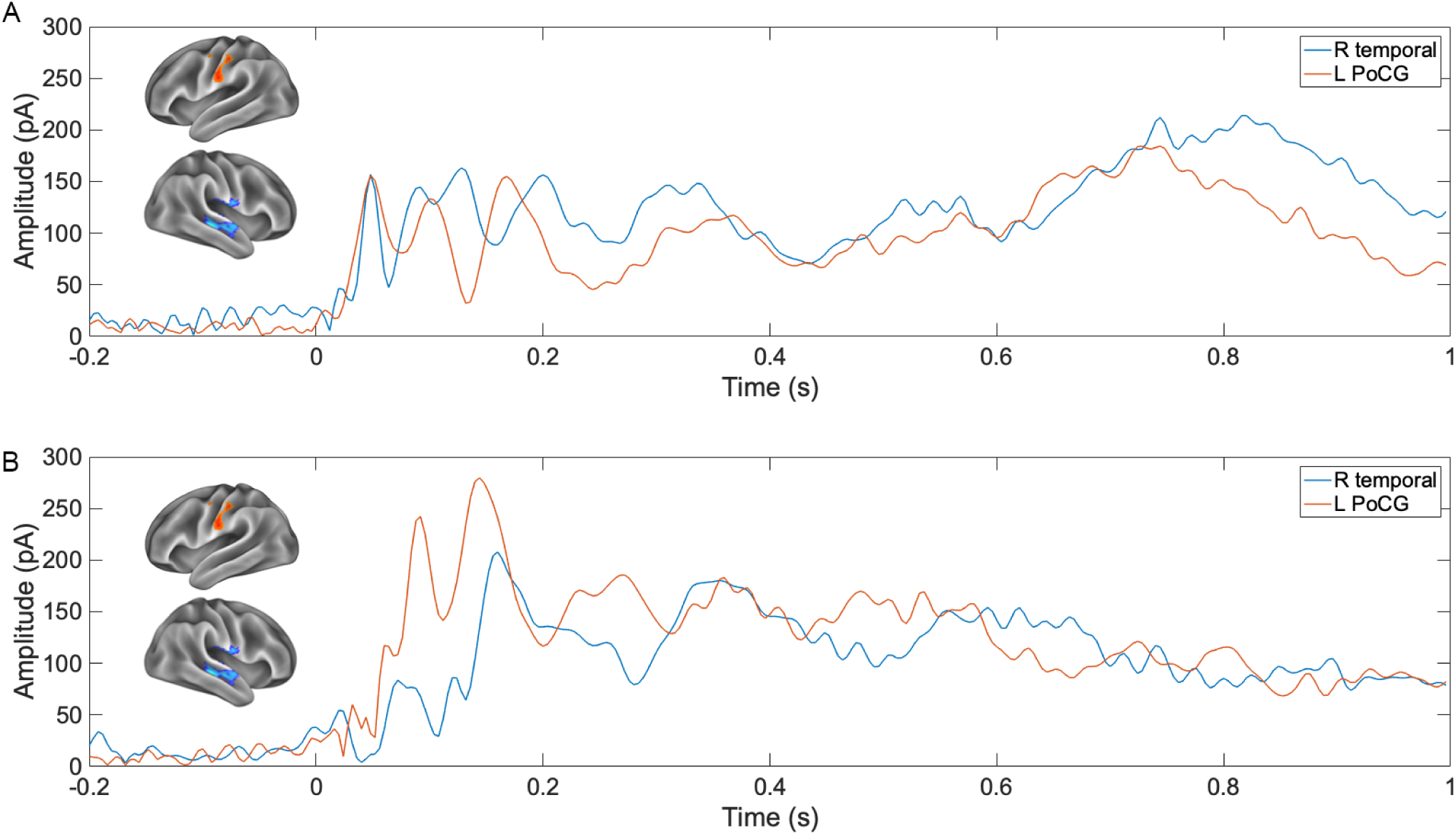
Source-estimated EEG activity within the right temporal and left post-central regions-of-interest identified in the fMRI scan. **(A)** Estimated activation in the ROI in response to auditory and **(B)** to vibrotactile versions of the same stimuli. Brain renderings show regions-of-interest used for source estimation.

## Discussion

We here, for the first time, demonstrate how training with a sensory substitution device (here, with a vibrotactile speech transducer) can recruit brain areas that are normally engaged by the processing of speech in the familiar (auditory) modality for the processing of speech in another (vibrotactile) modality. By training normally-hearing individuals on a set of vowel-consonant-vowel disyllables and acquiring fMRI-RSA scans before and after training, we showed that a network of regions across somatosensory, auditory, and multisensory association cortices represent VT speech after training. The left post-central gyrus in particular showed a training effect and increased its selectivity to VT speech after training. By acquiring an additional fMRI-RSA scan in which the acoustic versions of the VT speech stimuli were used, we showed that the right planum temporale is the one region that is selective for both acoustic and VT speech. Critically, the RSA provided evidence 1) for the recruitment of auditory speech regions by the processing of VT speech, and 2) for isomorphic representations between VT and auditory speech, thus supporting 3) the targeted “piggybacking” of the novel sense onto existing processing circuitry for the expert sense. Task-evoked functional connectivity analysis showed that after VT speech training, the left post-central gyrus was connected with bilateral superior temporal gyri and the right PT, suggesting that VT speech training resulted in the cross-modal recruitment of auditory areas. The temporal dynamics of VT speech perception were studied by projecting EEG data to source space. The left post-central gyrus was activated first, followed by the right temporal regions, suggesting that VT stimuli are first processed in left somatosensory regions before being relayed to right auditory areas.

Two brain regions of particular interest in the fMRI analysis were the left post-central gyrus and the right planum temporale. The left post-central gyrus was the one region that showed a significant increase in selectivity for VT speech after training. This region was functionally connected to regions within the auditory system after training, including the right PT and bilateral superior temporal gyri. The right PT was also selective for auditory speech, suggesting that this region is an interface point between the VT and auditory speech systems. The PT is a triangular region of auditory cortex posterior to Heschl’s gyrus and is a multifunctional nonprimary auditory region that is part of the early dorsal stream for the processing of complex spatio-temporal signals. There have been several postulated functions of the PT. (Griffiths and Warren, 2002) proposed that the PT is involved in the analysis of many types of complex sounds including speech and music, as well as the spatial and sensorimotor characteristics of sounds. According to this view, the PT is a “computational hub” that is involved in the segregation of incoming acoustic patterns and their spatial characteristics. The PT is also a critical part of the dorsal pathway in dual-stream models of auditory and speech processing (Hickok and Poeppel, 2007; Rauschecker and Scott, 2009). The dorsal stream is hypothesized to be involved in sensorimotor integration during speech processing, which has been described as an internal model process, a concept derived from motor control theory (Wolpert et al., 1995). For example, when producing speech, a forward model produces an “efference copy”, which is sent to sensory cortices to predict the upcoming sensory consequences of the motor command. An efference copy relayed to auditory cortices predicts what the utterance will sound like, and an efference copy sent to somatosensory cortices predicts the somatosensory and proprioceptive consequences of the utterance. The comparison of predicted (via efference copies) and actual sensory consequences is thought to be critical for vocal learning and the online control during speech production. The PT appears to be involved in this process. For example, in an fMRI study in human participants, produced speech was artificially modified in real-time by shifting the first speech formant frequency. The right PT was found to encode the mismatch between the expected and actual auditory signals (Tourville et al., 2008).

The PT has also been shown to respond to both visual and auditory linguistic stimuli, being activated by both spoken and written words (Nakada et al., 2001), as well as during silent lip-reading (Calvert et al., 1997). The PT also mediates the learned cross-modal correspondence of stimuli. During “key-touch reading,” a piece of music is identified by observing the pattern of piano key-touching movements. An fMRI study provided evidence that visual information during key-touch reading is transformed to the auditory modality through the PT (Hasegawa et al., 2004). An important open question is whether visual and somatosensory input to primary auditory cortex and PT is feedforward, or feedback from higher-level multisensory areas like the superior temporal sulcus. The opinion of (Calvert et al., 1999) was that enhanced activity in auditory cortex in response to audio-visual speech resulted from feedback. Further, electrophysiological experiments in non-human primates has demonstrated that visual stimulation results in feedback signals in auditory cortex (Schroeder and Foxe, 2002). The processing of somatosensory stimuli in auditory, on the other hand, is consistent with a feedforward pattern (Schroeder and Foxe, 2002; Schroeder et al., 2001). In our data, we find evidence for low-latency activity in right auditory areas after engagement of the left post-central gyrus, which is consistent with a feedforward model in which VT speech is first processed in the left post-central gyrus before being integrated with representations in right auditory regions. A possible function of this circuit is to map incoming VT input in somatosensory cortex onto existing auditory speech representations in auditory cortex.

An additional important question is how the functional connection between the left post-central gyrus and right PT is formed anatomically, and whether this connection is present prior to experience with VT speech and is “activated” by training. A DTI study provided evidence that the neural substrates of audition and somatosensation are linked anatomically between ipsilateral primary auditory and somatosensory cortices (Ro et al., 2013). The finding of extensive ipsilateral activity between the auditory and somatosensory cortices suggests that in our data the processing of VT speech might proceed first from the left to right post-central gyrus and then to the right auditory cortex. After nearly a century of research into using sensory substitution to convey speech information via the tactile sense, we here for the first time show a neural mechanism and spatiotemporal processing dynamics underlying VT speech learning. Our results provide evidence for a core circuit involving the left post-central gyrus and right planum temporale. The observation that both acoustic and VT speech co-localized to the PT, a key region in the dorsal stream thought to be critical, among others, for speech production (Chevillet et al., 2013; Hickok and Poeppel, 2004; Rauschecker and Scott, 2009) suggests that VT speech learning might benefit from training paradigms that couple perception and production. We also demonstrated isomorphic neural representations for VT and auditory speech (i.e., the representation for both modalities was organized according to phonological features). An interesting avenue for future investigation will be to determine whether isomorphic representations coupling VT and auditory speech processing at such low processing levels result in “automatic” processing of VT speech, and whether such a coupling scheme facilitates generalization to more complex, composite VT speech stimuli such as words, thought to be processed at higher levels of the auditory speech system.

## Acknowledgements

This work was supported by National Science Foundation Grant BCS-1439338 to M.R., National Science Foundation Grant BCS-1439339 to L.E.B., and National Institute on Deafness and Other Communication Disorders Award Number F30DC016496 to P.M. We thank Dr. Josef Rauschecker for helpful comments on this manuscript.

## Author contributions

P.S.M., S.P.E., E.T.A., L.E.B., and M.R. designed the experiment; P.S.M., S.P.E., and R.K. performed the experiment. P.S.M. analyzed the data. P.S.M., L.E.B., and M.R wrote the paper with assistance from S.P.E. and E.T.A.

## Declaration of interests

The authors declare no competing interest.

## Methods

### Participants

Twenty-seven right-handed healthy adults (ages 19-26 years, mean age=21.7 years, 17 females) were enrolled in the study. Georgetown University’s and George Washington University’s Institutional Review Boards approved all experimental procedures, and written informed consent was obtained from all participants before the experiment. Participants were paid for their participation.

### Vibrotactile speech display

A custom 17.4 cm x 11.0 cm 14-channel MRI-compatible vibrotactile stimulator array was organized as 2 rows of 7 stimulators (Fig. 1A), with center-to-center stimulator spacing of 2.54 cm. To ensure that the stimulators would maintain contact with the volar (i.e., same side as the palm of the hand) forearm, the array comprised four rigid modules connected with stiff plastic springs. Velcro straps were used to mount the device to the arm firmly while allowing the array to conform to the arm’s shape. The modules closest to the wrist (9.7 cm x 5.0 cm) each contained four stimulators, and the other two modules (7.1 cm x 5.0 cm) each had three stimulators. The piezoelectric bimorph stimulator wafers (www.piezo.com, model Q220-A4-303YB) were sandwiched between two custom-manufactured printed circuit boards (2-layer 1.5 mm FR-4 epoxy glass laminate) with 2.15-mm spacing between boards. Custom 3D-printed plastic contactors (with 4.6-mm diameters) were epoxied to the bimorph’s moving ends and protruded through 6.4-mm diameter surround holes in each circuit board. With no applied voltage to the piezoelectric bimorphs, the contactors were flush with the circuit board surface facing the skin. During operation, a constant +57-V voltage applied to all stimulators retracted the contactors into the surround, and each -85-V pulse drove the specified contactor into the skin. The drive signal was a square wave, with a pulse-time of 2 milliseconds (ms), and with unpowered intervals of 1ms between power reversals to protect the switching circuitry.

The display’s control system comprised the power supplies (-85V, +57V), high voltage switching circuits to apply the voltages to the piezoelectric bimorphs, and a digital control system that accepted from a controlling computer’s serial COM port the digital records specifying a stimulus (comprising the times and channels to output pulses on), and a command to initiate stimulus output.

### Vibrotactile speech stimuli

A real-time vocoder was used to convert recorded acoustic speech signals into vibrotactile stimuli. The initial stage of the vocoder comprised a bank of filters whose output power was used to control the output of vibrotactile pulses. The vibrotactile display used a frequency-to-place mapping algorithm: The energy passed by each filter of the vocoder was used to modulate the vibration of a specific transducer on the 14-channel VT device (Fig. 1A and 1B) placed on the volar forearm. Low frequencies mapped to transducers near the wrist, and higher frequencies mapped to transducers near the elbow. If the energy within a given filter exceeded a fixed threshold at a given time point, a vibrotactile pulse was emitted from the corresponding transducer. VT speech stimuli were generated using a range of thresholds, and a pilot experiment was performed to select a single threshold that resulted in maximally discriminable stimuli. The basic hardware design and software algorithms for the vocoder are referred to in (Bernstein et al., 1991) as the “GULin” vocoder algorithm, which had 16 channels. The ones used here were 13 bandpass filters (with center frequencies of 260, 392, 525, 660, 791, 925, 1060, 1225, 1390, 1590, 1820, 2080, and 2380 Hz (see the horizontal white lines in Fig. 1B), and with respective bandwidths of 115, 130, 130, 130, 130, 130, 145, 165,190, 220, 250, 290, and 330 Hz), and a 5000 Hz high-pass filter.

The stimuli were twelve recorded vowel-consonant-vowel (/a/-C-/a/, C=consonant) stimuli comprising the consonants /b, d, f, g, k, m, n, p, s, t, v, z/ with the vowel /a/. These stimuli had the same vowel, because training focused on distinctions of the phonological features of the consonant (see next section). Two tokens from the same male talker were used for each stimulus. To limit differences in amplitude that might be used as cues for discrimination, all of the auditory speech tokens were amplitude normalized to a total RMS of -14 decibels fullscale (dBfs) (i.e., the reference level for dBfs is defined as the maximum possible digital level) prior to conversion to the vibrotactile domain. The mean duration of the stimuli was 744 ms (range 665-830 ms). The mean consonant onset was 300.2 +/- 9 ms relative to stimulus onset. Consonant onset was determined by visual inspection of the acoustic waveforms and spectrograms.

### Training

Participants were trained to identify distinctions in phonological features, viz., voicing, manner, and place of articulation (Bernstein, unpublished experiments). The training task was to select which of two VT stimuli presented with an inter-stimulus-interval of 1000 ms matched an orthographic target (Fig. 1C) visually displayed on a screen. The two stimuli differed along a single phonological feature dimension (e.g., the two syllables differed in voicing but were matched for manner and place of articulation; see Table 1 for a list of all stimulus pairs). Participants responded with a button press to indicate which VT stimulus corresponded to the target. Participants received minimal feedback (“Correct!” or “Incorrect”) because minimal feedback preferentially engages an implicit procedural-based reflexive system that is beneficial for speech category learning (Chandrasekaran et al., 2014). During each training session, participants completed six blocks of 60 trials each. Each block was focused on training one of the three phonological features. To facilitate training progression, a weighting system was used, in which the stimuli that were mis-categorized most often were presented more frequently than ones that were correctly categorized: In each block, all stimuli were evenly distributed over 66% of the trials. The two stimuli that were most often mis-categorized from the previous block were used in the remaining 33% of trials. Participants were trained to 80% accuracy for each of the phonological feature contrasts to obtain a consistent level of performance across the sample of participants. Ear plugs and pink noise presented through ambient noise-dampening headphones were used to mask the sound of the VT device.

### Training analysis

Behavioral data from vibrotactile speech training were analyzed using generalized linear mixed-models (GLMM) in R using the glmer function from the lme4 package (Bates et al., 2015). Accuracy per-trial was modeled as the dependent variable using a logit link function, and training session and phonological feature contrast were modeled as fixed effects. For random effects, random slopes and intercepts were included in the model for each subject, and random intercepts were included for stimuli.

### Vibrotactile fMRI scan

A subset (n=20) of participants underwent vibrotactile fMRI scans both before and after VT speech training. EPI images from six short-block runs were collected. Each run lasted 7.6 minutes, beginning and ending with a 10-second fixation period. Within each run, VT stimulus repetitions were presented in 6-second blocks, with a 10-second inter-block interval. To boost the stimulus-driven BOLD signal, three repetitions of the same VT syllable were presented in each block. All twelve of the trained syllables were included in the scans. To maintain attention, participants performed an oddball detection task in the scanner. Participants pressed a button with their left hand whenever they felt an oddball stimulus. To generate a VT oddball stimulus, a modified vowel-consonant-vowel syllable was constructed, in which the consonant was replaced with broadband noise, and that stimulus was vocoded as described earlier. The oddball was presented in 28% of the blocks.

### Auditory fMRI scan

The same subset (n=20) of participants underwent an auditory fMRI scan after VT speech training and the post-training VT fMRI scan, in which they listened to the acoustic non-vocoded versions of the trained syllables. EPI images from six runs using a clustered-acquisition sequence were collected. Each run lasted 7.4 minutes and began and ended with a 20-second fixation period. Auditory stimuli were presented after every other volume, or every 6 seconds using Presentation (Neurobehavioral Systems) via customized STAX electrostatic earphones (https://staxaudio.com) at a comfortable listening volume (∼65-70 dB) worn inside ear protectors (Bilsom Thunder T1) giving ∼26 dB attenuation. Stimulus timing differed from the VT scan because of the use of a sparse-sampling design specialized for auditory experiments (Scheffler et al., 1998; Talavage and Hall, 2012). All twelve of the syllables in the training set were included in the scans. Participants performed an oddball detection task in the scanner. To generate an auditory oddball stimulus, a modified vowel-consonant-vowel syllable was constructed by replacing the consonant with an upwards tone-sweep. The tone-sweep was used rather than broadband noise (as used for the VT oddball stimulus), because during pilot testing it was discovered that the acoustic broadband noise resulted in the illusory percept of /asa/. Therefore, a tone-sweep was used, because it sounded sufficiently distinct from the trained syllables. Twenty-eight percent of the trials in the scan contained an oddball stimulus.

### MRI acquisition

MRI data were acquired at Georgetown University’s Center for Functional and Molecular Imaging using an EPI sequence on a 3-Tesla Siemens TIM Trio scanner. A 12-channel head coil was used (flip angle=90°, TE=29 ms, FOV=205 mm, 64×64 matrix). For the VT scans, 35 interleaved axial slices (thickness=4.0 mm, no gap; in-plane resolution=3.2×3.2mm^2^) were acquired with a continuous acquisition sequence (TR = 2040 ms). For the auditory scan, a clustered-acquisition sequence (TR = 3000 ms, TA = 1500 ms) was used such that each image was followed by an equal duration of silence before the next image was acquired. Twenty-eight descending axial slices (thickness=4.0 mm, no gap; in-plane resolution=3×3mm^2^) were acquired. A T1-weighted MPRAGE anatomical image (resolution 1×1×1mm^3^) was also acquired for each subject.

### fMRI data preprocessing

Image preprocessing was performed in SPM12 (http://www.fil.ion.ucl.ac.uk/spm/software/spm12/). The first four acquisitions of each run were discarded to allow for T1 stabilization, and the remaining EPI images were slice-time corrected to the middle slice (VT scan only; no slice-time correction was performed for the auditory scan due to its non-contiguous clustered-acquisition time series) and spatially realigned. EPI images for each subject were co-registered to their anatomical image. The anatomical image was then segmented, and the resulting deformation fields for spatial normalization were saved for later use when normalizing the RSA maps.

### Whole-brain searchlight representational similarity analysis (RSA)

RSA was used to localize the neural representations of VT and acoustic speech. In RSA (Kriegeskorte and Kievit, 2013), the dissimilarity of neural activation patterns elicited in response to different stimuli is compared to a posited model of the representational structure of those stimuli. A model of representation is tested by constructing a representational dissimilarity matrix (RDM), whose cells (row i, column j) correspond to the posited dissimilarity between stimulus *i* and stimulus *j*. We constructed a phonological feature RDM to investigate the localization of neural representations for VT and acoustic speech (Fig. 2A). Each entry of the phonological feature RDM quantified the dissimilarity between a pair of syllables along these three phonological feature dimensions (manner, place, voicing). For example, if a pair of syllables differed according to all three phonological features, the corresponding dissimilarity was one. If they differed according to one of the three phonological features, the dissimilarity was 0.33.

The CoSMoMVPA toolbox (Oosterhof et al., 2016) (http://www.cosmomvpa.org) and custom MATLAB code were used for the RSA. Each of the twelve syllables was modeled as a regressor in a first-level model. The onset of each trial was modeled using a canonical hemodynamic response function. Six motion parameters generated from realignment were included as regressors of no interest. *T*-statistic images were generated for the contrast of each stimulus condition (VCV syllable) relative to an implicit baseline. *T*-statistic maps were used, because *t-*values divide the *beta* estimate for each voxel by the estimate of its standard error, thereby reducing the influence of highly variable response estimates (Misaki et al., 2010). RSA was first performed on unsmoothed data and in participants’ native space, and then the RSA results for each individual subject were normalized to MNI space for statistical analysis. We performed a searchlight procedure (Kriegeskorte et al., 2006), in which the multivoxel response pattern associated with each speech stimulus was extracted from within a sphere of 30 voxels (similar results were obtained for a range of searchlight sizes from 20-100 voxels), and the dissimilarity between patterns for each stimulus pair was calculated using a correlation distance measure (1 minus the Pearson correlation). The mean of each feature (i.e., voxel) across conditions was subtracted prior to computing the dissimilarity for each stimulus pair (Diedrichsen and Kriegeskorte, 2017). The neural dissimilarity matrix for each searchlight was then Spearman-rank correlated to the phonological feature RDM, and the resulting correlation coefficient was assigned to the voxel at the center of the searchlight. This procedure was repeated for all searchlights across the entire brain, generating a whole-brain map of Spearman correlation coefficients between the neural dissimilarity matrix and the RDM. The resulting correlation coefficient maps were Fisher-*z*-transformed (atanh function in MATLAB) to conform to statistical assumptions for second-level parametric statistics. The Fisher-transformed maps for each subject were normalized to MNI space, smoothed with an isotropic 6-mm Gaussian kernel, and submitted to one-sample *t*-tests against zero using SPM’s second-level routines. Statistical maps were thresholded at a voxel-wise *p*<0.001 (uncorrected), and a cluster-level *p*<0.05 (FWE-corrected).

### fMRI functional connectivity

To characterize the connectivity of brain regions of interest identified in the RSA, seed-to-voxel functional connectivity was analyzed using the CONN-fMRI toolbox (Schurz et al., 2015; Whitfield-Gabrieli and Nieto-Castanon, 2012). Functional images were normalized into MNI space, and *beta* images were generated for all non-oddball stimuli. Noise due to white matter and CSF signals was regressed out using CompCor (Behzadi et al., 2007). Movement parameters were entered as covariates of no interest. Main condition effects (block onsets and durations) were also included as covariates of no interest to ensure that temporal correlations between BOLD time courses reflected functional connectivity and did not simply reflect stimulus-related coactivation. The BOLD signal time series from a region of interest was extracted for each subject and correlated with the time series of every voxel in the brain to generate a functional connectivity map. Individual subject connectivity maps were entered into a second-level random-effects analysis to assess connectivity at the group level. Results were thresholded at a voxel-wise *p*<0.001 (uncorrected), and a cluster-level *p*<0.05 (FWE-corrected).

### EEG experiment

A subset of participants (n=18) underwent an EEG experiment at George Washington University after VT speech training and after the completion of all fMRI experiments. Twelve of these participants also underwent fMRI. Participants carried out a VT oddball detection task. The same VT stimuli (trained syllables and oddball) were used in the fMRI and EEG experiments. Participants completed 3 blocks of 213 trials each using a pseudorandomly jittered inter-trial-interval of 1-1.5 s. The oddball was presented pseudorandomly on 28% of the trials.

### EEG acquisition

EEG recording used a 64-electrode Acticap with active Ag/AgCl electrodes. Bipolar EOG electrodes were affixed around the eyes to monitor eye movements. The EEG was amplified using a high dynamic range amplifier (SynAmps 2, Neuroscan) and digitized at 1000 Hz with a 200-Hz low-pass filter. Electrode impedances were measured with an inclusion criterion of 10 kOhm. Electrode position was registered using a Polhemus FASTRAK.

### EEG preprocessing

Offline, the data were highpass filtered at 0.1 Hz with a 24- dB/octave roll off FIR zero phase-shift filter using Edit 4.5 software (Neuroscan, NC). To reduce the presence of eyeblink artifact in the data, the peak of eyeblink activity was identified in the vertical eye movement bipolar channel in the continuous data with a simple voltage-triggering algorithm. The continuous data were epoched around the detected eyeblinks, the morphology of the eyeblink artifact was reviewed visually for the presence of other artifacts, and the remaining eyeblinks were then averaged for each subject. Within the Edit software, a spatial singular value deconvolution (Spatial SVD) was then used to generate a set of coefficients characterizing the distribution and time– amplitude function for the eyeblink artifact. A spatial filter was generated in Edit using the coefficients and then applied to the filtered continuous data file, substantially reducing the contamination of the data. Each participant’s data were then imported into EEGLAB ((Delorme and Makeig, 2004) with electrode positions recorded using Polhemus. The data were resampled to 500 Hz. EEGLAB’s downsampling function employs a zero-phase anti-aliasing filter prior to downsampling. Bad channels (flatline or noisy channels, or channels with low-frequency drifts) were identified, removed, and interpolated using the clean_rawdata plugin in EEGLAB (on average 7 +/- 1.4 channels per subject). Data were then epoched from -0.2 to 1 second relative to stimulus onset. Epochs with peak-to-peak amplitude fluctuations of greater than 150 microvolts were automatically rejected (on average 116 +/- 36.7 epochs per subject). Next, data were average re-referenced and baseline corrected (baseline defined as -0.2 to -0.1 seconds relative to stimulus onset). The EEG data were submitted to independent component analysis (ICA) using the infomax ICA algorithm in EEGLAB to identify and remove artifacts such as artifacts associated with ocular movement, other muscle movement, and VT stimulator artifacts. The SASICA (Semi-Automated Selection of Independent Components) toolbox was used to aid in the selection of artifactual components (Chaumon et al., 2015). ICA was performed separately for VT and auditory experimental blocks. Components marked as artifactual were removed from the data. On average, 13.2 +/- 1.0 components out of 64 total components were rejected for each subject for the auditory experimental blocks, and 12.2 +/- 0.8 components out of 64 total components were rejected for each subject for the VT experimental blocks. Finally, the ICA-denoised data were low-pass filtered at 40 Hz and resampled to 250 Hz. Single trial event-related potential (ERP) estimates for each VT syllable condition were then averaged in preparation for RSA.

### EEG representational similarity analysis

In order to determine the temporal dynamics of phonological feature selectivity during the processing of VT and acoustic stimuli, EEG RSA was performed using the CoSMoMVPA toolbox (Oosterhof et al., 2016). A searchlight algorithm with a sliding time-window was implemented in which RSA was repeated in circumscribed spatial neighborhoods of electrodes (*n*=6) across time using a sliding time-window of 28 ms (similar results were obtained for other choices of neighborhood and window size). The analysis was replicated for searchlight sizes of 3-8 electrodes and for time windows of 15-30 ms and the results were qualitatively consistent (data not shown). At each time window and searchlight cluster, a neural RDM was computed by taking the pairwise dissimilarity between the ERPs for each stimulus using the correlation distance measure. The resulting neural RDM was then Spearman correlated with the phonological feature RDM, and the resulting correlation coefficient was assigned to the corresponding time point and electrode. Statistical testing to identify spatiotemporal electrode clusters with correlations significantly above zero was performed using a threshold-free cluster-estimation procedure (Smith and Nichols, 2009) using multiple-comparison correction based on a sign-permutation test as implemented in CoSMoMVPA.

### EEG source estimation

EEG analysis was performed in source space in order to assess the temporal dynamics of activation within regions-of-interest involved in the representation of VT speech identified in the fMRI. Cortical source activations were estimated using the Brainstorm toolbox (Tadel et al., 2011). Current sources were approximated using the standardized low-resolution electromagnetic topography algorithm (sLORETA) (Pascual-Marqui, 2002). The EEGLAB preprocessed data were imported into Brainstorm. Pre-stimulus baseline intervals (-200 to -100 ms) were used to calculate single subject noise covariance matrices. The forward model that characterizes the contribution of source activity to EEG activity at the scalp was computed using the Boundary Element Method (BEM) as implemented in OpenMEEG (Gramfort et al., 2011). The option of constrained dipole orientations was selected for source estimation. To estimate the temporal dynamics of activity within ROI identified in the fMRI-RSA, masks of the fMRI ROI were imported into Brainstorm and projected into source space.

## Notes

### Competing Interest Statement

The authors have declared no competing interest.

